# A novel localization based biosensor, SEDAR, visualizes spatio-temporal dynamics of activated Ras in endogenous *Drosophila* tissues

**DOI:** 10.1101/432997

**Authors:** Rishita Changede

## Abstract

Chemokine signaling via growth factor receptor tyrosine kinases (RTKs) regulates development, differentiation, growth and disease implying that it is involved in a myriad of cellular processes. A single RTK, for example the Epidermal Growth Factor Receptor (EGFR), is used repeatedly for a multitude of developmental programs. Quantitative differences in magnitude and duration of RTK signaling can bring about different signaling outcomes. Understanding this complex RTK signals requires real time visualization of the signal. To visualize spatio-temporal signaling dynamics, a biosensor called <u>SE</u>nsitive <u>D</u>etection of <u>A</u>ctivated <u>R</u>as (SEDAR) was developed. It is a localization-based sensor that binds to activated Ras directly downstream of the endogenous activated RTKs. This binding was reversible and SEDAR expression did not cause any detectable perturbation of the endogenous signal. Using SEDAR, endogenous guidance signaling was visualized during RTK mediated chemotaxis of border cells in *Drosophila* ovary. SEDAR localized to both the leading and rear end of the cluster but polarized at the leading edge of the cluster. Perturbation of RTKs that led to delays in forward migration of the cluster correlated with loss of SEDAR polarization in the cluster. Gliding or tumbling behavior of border cells was a directly related to the high or low magnitude of SEDAR polarization respectively, in the leading cell showing that signal polarization at the plasma membrane provided information for the migratory behavior. Further, SEDAR localization to the plasma membrane detected EGFR mediated signaling in five distinct developmental contexts. Hence SEDAR, a novel biosensor could be used as a valuable tool to study the dynamics of endogenous Ras activation in real time downstream of RTKs, in three-dimensional tissues, at an unprecedented spatial and temporal resolution.

## Significance

Chemokine signaling is vital for many cellular processes. It is thought that the strength and duration of the RTK signaling can regulate different downstream outcomes in invivo tissues. A biosensor is needed to visualize these quantitative differences. SEDAR, a biosensor was developed to visualize Ras activation immediately downstream of the RTKs. This sensor visualized activation of endogenous receptors in real time, and subcellular spatial resolution. SEDAR visualized signaling in several tissues, in Drosophila and can be used to understand signaling in various contexts.

## Introduction

Growth factor receptor tyrosine kinases (RTKs) are chemokine receptors that signal during key developmental processes (1–3) and chemotaxis (4). Growth factor receptor such as EGFR are often mutated in several cancers and are indicators of poor prognosis (5). The myriad of outcomes of EGFR are brought about by few intracellular signaling pathways (6, 7). One way to bring about different signaling outcomes is to regulate the magnitude i.e. strength and duration of RTK activation (8, 9). Hence, visualization of endogenous RTKs signaling in real time and at subcellular resolution could provide critical insights into the diverse outcomes downstream of RTKs.

Forster Resonance Energy Transfer (FRET) based biosensors such as Raichu, reported that dynamics of Ras downstream of growth factor receptor signaling (10, 11). However FRET probes are not used *in vivo* to visualize dynamics of endogenous RTK signaling due to poor technical challenges such as poor FRET efficiency deep within the tissue (12) and dominant negative effects of high expressing FRET constructs that disrupt very subtle endogenous RTK signals (13–15). RTK signaling is typically visualized *in vivo* using reporter constructs expressing GFP/LacZ or phospho specific antibodies to downstream components such as Erk are suitable to provide binary information of RTK activation (3, 16). To date, following the dynamics of this activation in real time, that are instructive for the cell, is not possible (8). Direct visualization of *in vivo* RTK signals pose problems in that, they are very subtle and antibodies to phosphoPVR could not detect the subtle activation of PVR without amplification by overexpressing PVR (15). Recent approaches use phase separation based *in vivo* assays to visualize dynamics of Erk signaling (17) which is several steps downstream of the RTK signaling cascade and does not provide information on RTK activation dynamics. Mass spectrometry based assay of phosphorylation events downstream of EGFR activation were visualized at seconds’ time scale (18), yet these may be challenging experiments to perform *in vivo*. Hence a biosensor to visualize endogenous RTK activation dynamics at subcellular resolution is much needed.

An effective biosensor to sense endogenous signals would need very low transgene expression, both to detect subtle signals and to avoid interference with endogenous signaling while maintaining high sensitivity towards the signal. This paper reports the development of a novel localization based biosensor probe, SEnsitive Detection of Activated Ras (SEDAR), to visualize the spatio-temporal dynamics of endogenous RTK signals in *in vivo*. Activation of Ras, a small GTPase downstream of RTKs (19) and upstream of multiple signaling pathways (20) was visualized by binding of highly fluorescent Ras binding domain of Raf to it (See Figure 1). SEDAR visualized signaling dynamics in RTK guided (PVR and EGFR) chemotactic border cell migration that occurs during *Drosophila* oogenesis (16, 21, 22). Border cells are a group of 6-8 cells that delaminate from the follicular epithelium and migrate through the germline towards the developing oocyte (23, 24). SEDAR polarized at the leading edge of the cluster and the magnitude of this polarization determined the size of protrusions. In addition to border cells, SEDAR detected the subcellular localization of activated Ras immediately downstream of EGFR signaling in several tissues during *Drosophila* development (1, 3). Hence SEDAR can have wide spectrum of applications in understanding the complex regulation of varied signals mediated by Ras activation. Taken together, SEDAR is a novel biosensor that visualized spatio-temporal dynamics of Ras activation in *in vivo* tissues.

**Figure 1:**
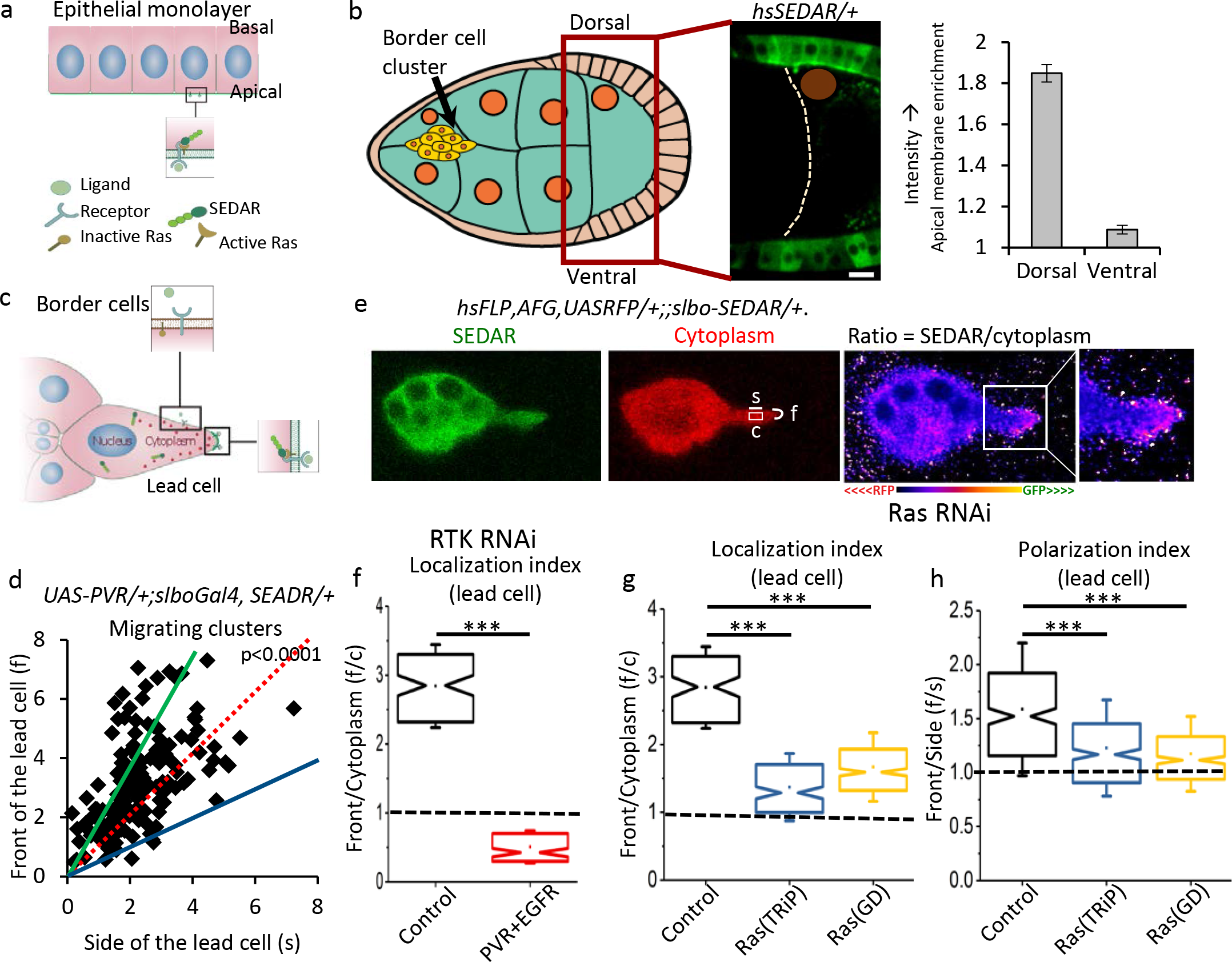
SEDAR visualized RTK activation downstream of Ras signaling (a) schematic representation of localization of SEDAR to the apical membrane upon activation of RTK signal in epithelium. (b) schematic representation of Drosophila egg chamber where the Dorsal follicular epithelium cells (in proximity to the oocyte nucleus) respond to Gurken signaling from the oocyte. Red box marked the region shown in SEDAR panel. SEDAR panel showed localization of SEDAR in the follicular epithelium, orange circle-oocyte nucleus. Graph quantified the membrane localization of SEDAR normalized to cytoplasm in the dorsal and the ventral follicle cells(n=46, 12 egg chambers) (c) schematic of SEDAR localization in the lead cell of the border cell cluster in response to RTK signaling (d) Scatter plot of intensity at the front (f) versus the side (s) of the leading cell in wild type early migrating border cell clusters, f=2s green line, f=s red line, 2f=s blue line, n=166 distinct clusters, p value computed by Wilcoxon singed test (e) control: *hsFLP,AFG,UAS-RFP/+; SEDAR/+*. Ratiometric image of SEDAR over cytoplasm in pseudo color (LUT-fire). Box marked the zoom in region shown in the last panel. ROI measured in the ration image were marked in red channel. (f & g) Box plot of localization index of the lead cell of the border cell cluster in Control and RTKS RNAi (PVR and EGFR, f): *hsFLP,AFG,UAS-RFP/hsFLP; UAS-EGFR-RNAi/+;SEDAR/UAS-PVR-RNAi* or RasRNAi: *hsFLP,AFG,UASRFP/hsFLP;;SEDAR/UAS-RasRNAi* (h) Box plot of Polarization index of lead cell of the border cell cluster in control and RasRNAi. In all box plots, the box represented the upper and lower quartile, the notch represented the median, the dot represented the mean, whiskers represent standard deviation and dotted line marked equal ratio. Control n= 266, 5 movies; RTKRNAi n=180, 4 movies, Ras(TRiP) RNAi n=315, 6movies, Ras (GD) RNAi n=320, 6movies. ***p<0.0001, Scale 10µm.

## Results

### Design and validation of the biosensor

Ras, a GTPase activated downstream of several RTKs (for example, PVR, EGFR, (16, 25–27) was visualized by specific recruitment of Ras binding domain (RBD) of Raf, a MAPKKK, to GTP bound activated Ras (Figure S1a). Different combinations of minimal or extended RBD were fused to a linear string of ten molecules of GFP (Figure S1b). Combined fluorescence of ten GFP molecules increased the sensitivity of visualization of the RTK signal and allowed for low expression of the transgene. For ubiquitous expression, a gentle and acute induction of a heat-shock promoter (hs) was used (see methods). Expression under a UAS promoter gave rise to high expression, inadequate to detect low signals.

To identify the most suited biosensor design, localization of all hs-sensors were observed in a well-documented example of RTK signaling in the *Drosophila* ovary, namely EGFR signaling in response to the dorsal ventral graded distribution of Gurken ligand secreted by *Drosophila* oocyte. Dorsal follicular epithelium activated EGFR signaling whereas non signaling ventral cells serve as a negative control (28, 29) (Figure 1a, 1b). The signal was quantified as intensity of GFP at the apical plasma membrane of follicle cells normalized to the cytoplasmic intensity of GFP (Figure 1a, 1b). Several Ras biosensors did not localize specifically to the plasma of the dorsal follicular epithelium (Figure S1b, S1c). Two of the five RAS biosensors (HS-RBD-10XGFP and HS-2RBD-10xGFP) were preferentially recruited to the apical plasma membrane in the dorsal cells, with no detectable recruitment in ventral cells (Figure 1b, S1b, S1c). Sensitivity of 2RBD-10xGFP was higher than RBD-10XGFP (Figure S2a) and was chosen for further experiments and it was called SEDAR.

To validate SEDAR as a sensor for activated RTKs, we tested SEDAR localization in border cell clusters overexpressing PVR. Janssens et al, showed preferential enrichment of phosphoPVR to the front of these clusters. SEDAR enrichment was distinctly visible in epithelium where apical plasma membrane had very regular boundaries, however such distinct enrichment was not immediately visible in border cell clusters. To measure subcellular localization of SEDAR in border clusters over expressing PVR, a region of interest were marked by line 0.4µm thick and 1-3μm long at the front edge(f) and side edge(s) of the same cell in image of the membrane dye and the average intensity measured in the SEDAR image (Figure S2a). SEDAR mimicked phosphoPVR localization and showed a preferential enrichment at the front compared to the side of the leading cell in the border cell cluster (Figure 1e, n=166 clusters, Figure S2a) suggesting that SEDAR could sense RTK signaling.

### A ratiometric method for robust detection of SEDAR in cells with irregular membrane

Membrane label marked the plasma membrane of the border cell cluster and the nurse cells, introducing errors in the measurements hence neutral cytoplasmic marker, slbo-TurboRFP, was co-expressed to demarcate the border cell clusters. Due to the polyploid nature of the border cells, the expression pattern of SEDAR in different border cells varied (Figure S1d) so mild expression of SEDAR was induced specifically in the border cells under the control of the genomic region upstream of *slbo* and its 5’UTR (Figure 1d). A ratio of SEDAR image to the cytoplasmic RFP image (Figure 1e, Ratio) showed preferential SEDAR enrichment at the front of the border cell cluster (Figure 1e, last panel), making it well suited to measure plasma membrane recruitment of SEDAR in cells with irregular boundaries.

To measure subcellular localization of SEDAR, ROIs were marked in the RFP image, at the front (f) and side (s) of the leading cell, and measured in the Ratio image (Figure 1e). All measurements were normalized to cytoplasm (box‘c’, Figure 1e), marked to carefully avoid the nucleus Figure1d). This parameter was termed localization index (f/c). Value of 1 or less corresponded to no enrichment of SEDAR at the plasma membrane, greater than 1 indicated recruitment of SEDAR at the plasma membrane. To test if this was a general enrichment of SEDAR at the entire plasma membrane or a specific enrichment at the front edge of the leading cell, the signal at the front edge and side were compared (f/s). This was termed as the polarization index. Wild type clusters had a high localization (2.84±0.59, Mean±SD, Figure 1e, 1f - control) and polarization index (1.59±0.57, Figure 1e, 1h-control). phosphoPVR antibody could only detect signal upon PVR overexpression and was not sensitive enough to report the subtle endogenous signal. This is the first observation of the endogenous RTK signaling. Migration speed of the border cell clusters is sensitive to perturbation of RTK signal (26). Expressing of SEDAR did not perturb this migration speed (Figure S2b) showing that low levels of endogenous guidance signal could be monitored in cells with irregular and dynamically changing cells without disrupting it.

### SEDAR localization was stringently dependent upon RTK and Ras activation

To test the specificity of SEDAR signal to guidance input provided by PVR and EGFR, both receptors were simultaneously knocked down using RNAi (Figure 1f, (30). Expression of RNAi disrupted SEDAR recruitment to the plasma membrane (Figure S2c). SEDAR localization index was reduced below 1. As there was no preferential recruitment at the plasma membrane, if the measurement method is accurate the polarization index should also be below 1. It was measured to be 0.93 (±0.27). This result confirmed that SEDAR localization at the leading edge of border cell cluster stringently depended on guidance signal provided by the RTKs. Thus, SEDAR specifically detected subcellular signaling from endogenous guidance receptors during invasive cell migration.

To validate that SEDAR detected signaling by Ras, two distinct RNAi construct targeting the major Ras, Ras85D, were expressed individually with SEDAR. SEDAR intensity at the membrane was not visible in RNAi expressing border cell clusters (Figure S2c). Ras RNAi expression decreased localization index to 1.66 (±0.5) in GD line and 1.37 (±0.51) in TRiP line (Figure 1g), possible due to incomplete knockdown of Ras. The polarization index was 1.17 (±0.35) and 1.22(±0.41) respectively in the two RNAi lines (Figure 1h). In contrast, expression of RasV12, a constitutively activated form of Ras, showed an enrichment of SEDAR at the entire plasma membrane of all border cells (Figure S2e). Together, it showed that SEDAR localization at the leading edge specifically reported activation of Ras.

### SEDAR visualized dynamics of RTK signaling during border cell migration

We next assayed SEDAR localization in real time. It showed a front localization compared to the side of the same cell (Figure 2a, 2b). Average signal over the entire outer membrane of the border cell cluster was much lower and the bias was averaged out, serving as a control (Figure 2b). SEDAR localization at the leading edge of the cluster showed a cyclic pattern of increasing and decreasing signal, which was not observed at the side of the cell (Figure 2a, 2b, Video S1), similar to that observed in migratory cells in tissue culture (31).

**Figure 2:**
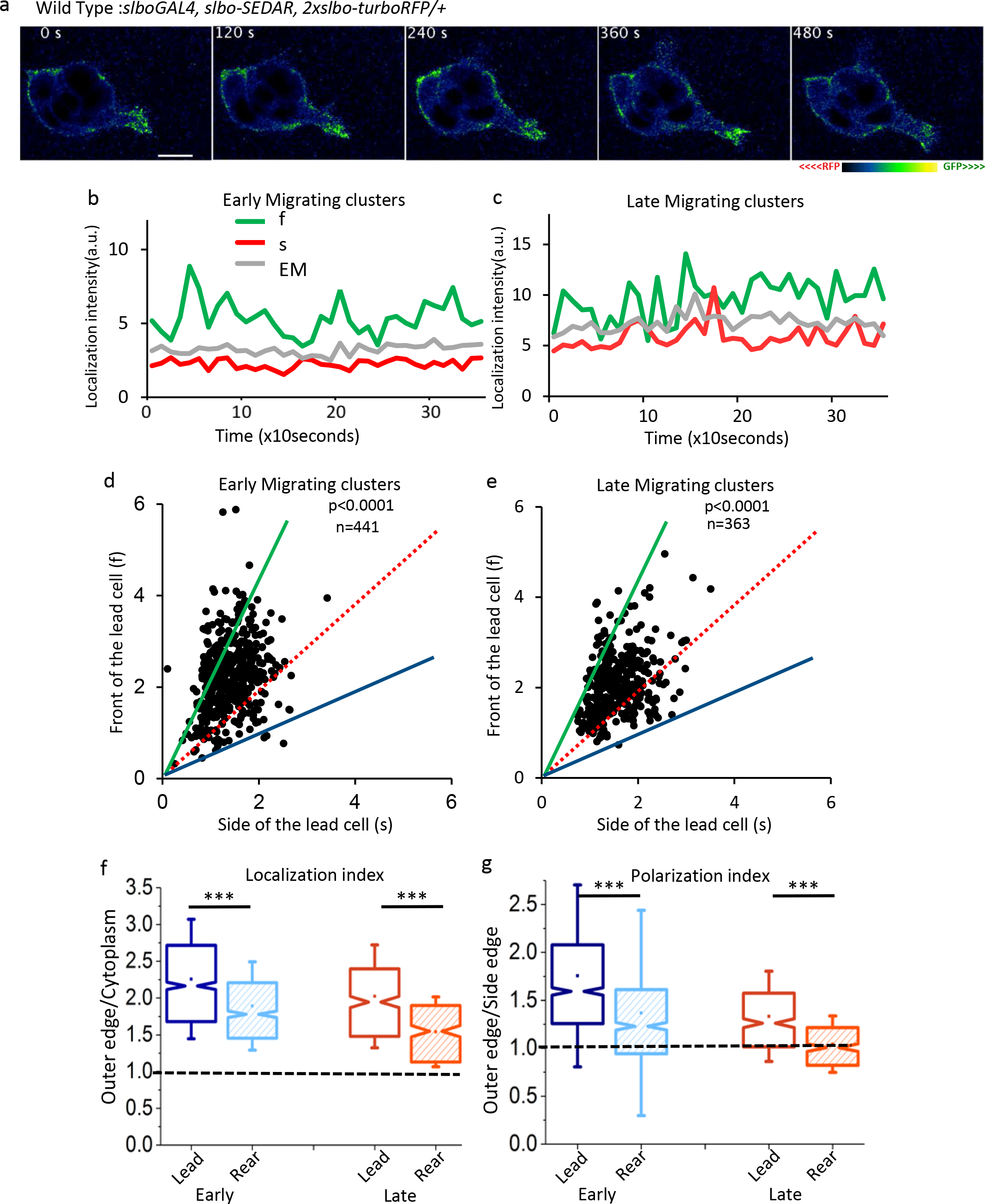
SEDAR localization was polarized within the lead border cell and the border cell clusters. (a) Ratiometric image sequence of SEDAR/neutral cytoplasm) of wild type early migratory cluster. Pseudo color (green fire blue) represented the ratiometric image, as indicated below. Scale −10μm. (b&c) Mean intensity of front (f, green), side (s, red) of the lead cell and outer membrane of the entire cluster (EM, grey) over time in a migrating border cell cluster. (d&e) Scatter plot of mean intensity of front(f) and side(s) membrane of the lead cell in early and late migratory cluster respectively. f=2s green line, f=s red line, 2f=s blue line, p by Wilcoxon singed test (f&g) Graph of localization index and polarization index of lead cell and rear cell in early migratory and late migratory wild type border cell clusters. In all box plots, the box represented the upper and lower quartile, the notch represented the median, the dot represented the mean, whiskers represent standard deviation and dotted line marked equal ratio. In early migratory clusters, Lead cell - n=1049, 20 movies; rear cell-n=710, 12 movies. Late, lead cell, n=360, 6 movies, rear cell – 240, 5 movies.

We assayed whether border cell behavior was directed by spatial activation of RTK within the border cell clusters. Border cell migration was divided into early phase, until the border cells reach 50% distance during the posterior migration and the late phase from 50% until they reach the oocyte (30). Early migration is characterized by persistent gliding movement with a dominant leader cell that has large protrusions whereas the late phase of migration is characterized by tumbling behavior where leader cells are frequently exchanged and have short lived, small protrusions. Strong front bias was observed in the leading cell of early migratory clusters (Figure 2d). Front bias was also observed in late migratory cluster (Figure 2c, 2e)

Localization and polarization index of SEDAR were measured in the lead and the rear cell. For the rear cell, these parameters were measured at the back of the rear cell (‘r’ Figure S3a) compared to its cytoplasm or side of the same cell. Polarization index of front cell over the rear cell measured the cell extrinsic cluster polarity of RTK activation.

In wild type early migratory clusters, the leading cell showed a higher localization index (2.2 ±0.81, Figure 2f) compared to the rear cell (1.89±0.6, Figure 2f) suggesting an overall front bias of SEDAR localization in the migrating border cell cluster. In the front cell, polarization index was higher (1.75±0.96, Figure 2g) compared to the rear cell (1.37±1.07, Figure 2g). Cell extrinsic cluster polarization was 28.5 percent higher at the front compared to the back of the entire cluster. In late migratory clusters, the localization index was 2.02 (±0.7) in the lead cell and 1.54 (±0.47) in the rear cell (Figure 2f). It is reduced compared to the early clusters but there is signal. The polarization index was reduced to 1.33 (±0.47) in the lead cell and 1.04 (±0.29) in the rear cell (Figure 2g). The magnitude of the cell intrinsic polarity in the lead cell was reduced yet the cell extrinsic cluster polarity was maintained at 27.9 percent higher in the front of the cluster compared the back of the cluster. This indicated that cell extrinsic cluster polarity dictated forward migration and magnitude of cell intrinsic polarity could dictate the mode of migration i.e. gliding or tumbling. Low magnitude of outward polarization in the rear cell correlated with fewer protrusions observed in the rear cell (30). SEDAR was assayed upon RTK perturbation which mimicked specific aspects of directed migration or migratory behavior

### Spatial localization of RTK activation determined behavior and migration of border cells

RTK perturbation disrupted directed migration or (overexpression) mimicked specific aspects of early or late migratory behavior. For these analyses, early migratory clusters were imaged. This allowed for comparison of clusters at the similar stage of migration cells as PVR^DN^EGFR^DN^ and EGFR overexpressing clusters were delayed during migration (30).

To test if intracellular polarization index of SEDAR correlated with directed migration, SEDAR localization was assayed in border cells clusters where directed migration of the cluster was disrupted by simultaneous overexpression of the dominant negative RTKs (RTK^DN^-EGFR^DN^PVR^DN^) without affecting the motility of the individual cells (30). Over time, no preferential front localization of SEDAR was observed at the front of the leading cell (Figure 3a, 3b). Though SEDAR localized to the plasma membrane (Figure 3c, Table S1). It was diffused as observed by the polarization index value of ^~^1 (lead cell – 1.16±0.44, rear cell-1.2±0.43, Figure 3d, Table S1) indicating loss of cell intrinsic polarity. The cell extrinsic polarization of the border cell cluster was lost with a marginal increase in the rear cell (3.2 percent) compared to the lead cell. This could explain the loss of directed migration observed within these clusters (30). SEDAR localization showed that cell extrinsic polarization of RTK signal in the border cell cluster guided migration of it. Further, no persistent protrusion was observed in the lead cell correlating with low magnitude of RTK polarization in the lead cell.

**Figure 3:**
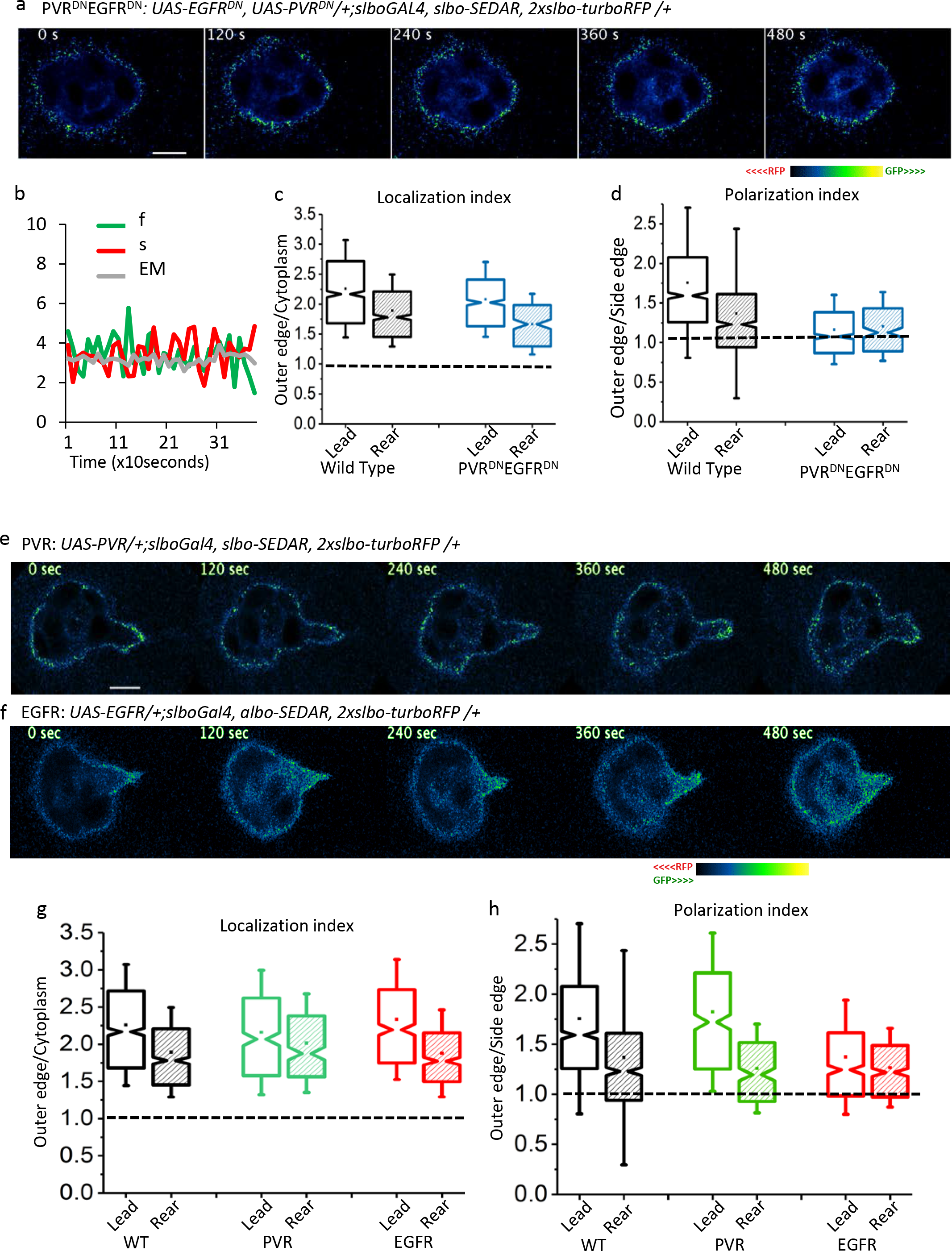
SEDAR polarization within the border cells correlated directed behavior of border cell clusters (a, e & f) Ratiometric image sequence of SEDAR and RFP Cytoplasm of genotypes indicated above the images. Scale 10μm (b) Graph showing time fluctuations of the intensity of front of leading cell, side of leading cell and entire outer membrane of the border cell cluster (c & g) Box plot of localization index or (d & h) polarization index in lead and rear cell of border cell cluster. In all box plots, the box represented the upper and lower quartile, the notch represented the median, the dot represented the mean, whiskers represent standard deviation and dotted line marked equal ratio. Wild type lead-n=1049, 20 movies, rear-n=710, 12 movies; PVR^DN^EGFR^DN^ lead-n=744, 13 movies, rear-n=240, 4 movies, PVR lead-n=379, 9 movies, rear-n=217, 4 movies, EGFR lead-n=425, 9 movies, rear-n=420, 7movies.

PVR overexpression promoted efficient gliding movement typical of the early phase of migration (26, 30). In border cell clusters overexpressing PVR (Figure 3e), the polarization index of SEDAR in the lead cell is increased to 1.82 (±0.79, Figure 3h), yet the amount of signal within the leading cell is comparable to wild type (localization index lead cell – 2.15±0.83, rear cell – 2.01±0.66, Figure 3g, Table S1). This correlated well with the persistent leader cell and large strong protrusions. In the rear cell, SEDAR localization is outward polarized 1.26 (±0.44) but reduced slightly compared to the wild type clusters (Figure 3h). The cell extrinsic cluster polarity is increased to 44.6 percent compared to the 28.3 percent in the wild type. This correlated well with persistent forward migration of the border cell cluster.

EGFR overexpression operated primarily via tumbling mode of migration where a persistent leader cell was absent (26, 30). In border cell clusters overexpressing EGFR (Figure 3f), SEDAR showed comparable localization index (lead cell-2.33±0.8, rear cell 1.88±0.58, Figure 3g). However, the polarization index of the lead cell was reduced to 1.37 (±0.57, Figure 3h) comparable to the late phase of migration. Next, the rear cell also showed an outward polarization of 1.26 (±0.39, Figure 3h) which reduced the cell extrinsic cluster polarity to a mere 8.3 percent; over five-fold lower than that observed within the clusters overexpressing PVR and three-fold lower than wild type early clusters. Loss of cell extrinsic cluster polarity correlated with severe delays in migration of the border cell clusters as overserved with EGFR overexpression (reported earlier, (30), suggesting that it determined forward movement of the cluster. Hence, SEDAR localization patterns detected activation of both PVR and EGFR.

Low magnitude of cell intrinsic polarity of SEDAR recruitment in lead cell of late migratory cluster, rear cells of early and late migratory cluster, and lead cell of EGFR^DN^PVR^DN^and EGFR overexpression explained the observed absence of large protrusions in each of these cluster (30) whereas large protrusions observed in lead cell of early migratory cluster and PVR overexpression correlated with large protrusions and gliding motility (30). Next, high cell extrinsic cluster polarity determined effective forward migration without delays as observed in PVR overexpression, WT early and late migratory clusters and reduction in this polarity led to severe delays as observed in PVR^DN^EGFR^DN^and EGFR overexpression (30). Taken together, these results showed that differential spatial activation of RTK signaling at the plasma membrane could directly alter behavior and migration of border cells.

### SEDAR localized to plasma membrane downstream of EGFR or PVR signaling in different tissues in *Drosophila*

Could SEDAR sense RTK activation in different tissues? EGFR signaling is key for many events during development. SEDAR localization was assayed in tissues which use EGFR signaling during development.

During *Drosophila* embryogenesis, EGFR function is required in the epidermis for proper dorsal closure and its activity is observed by phosphoErk (32). SEDAR was recruited to the plasma membrane of the epidermis during dorsal closure (Figure 4a). Stage 16 embryos served as negative control, where no EGFR signaling is reported in the epidermis. In these embryos, SEDAR appeared diffused in the cytoplasm and no preferential localization was observed at the plasma membrane (Figure S4).

**Figure 4:**
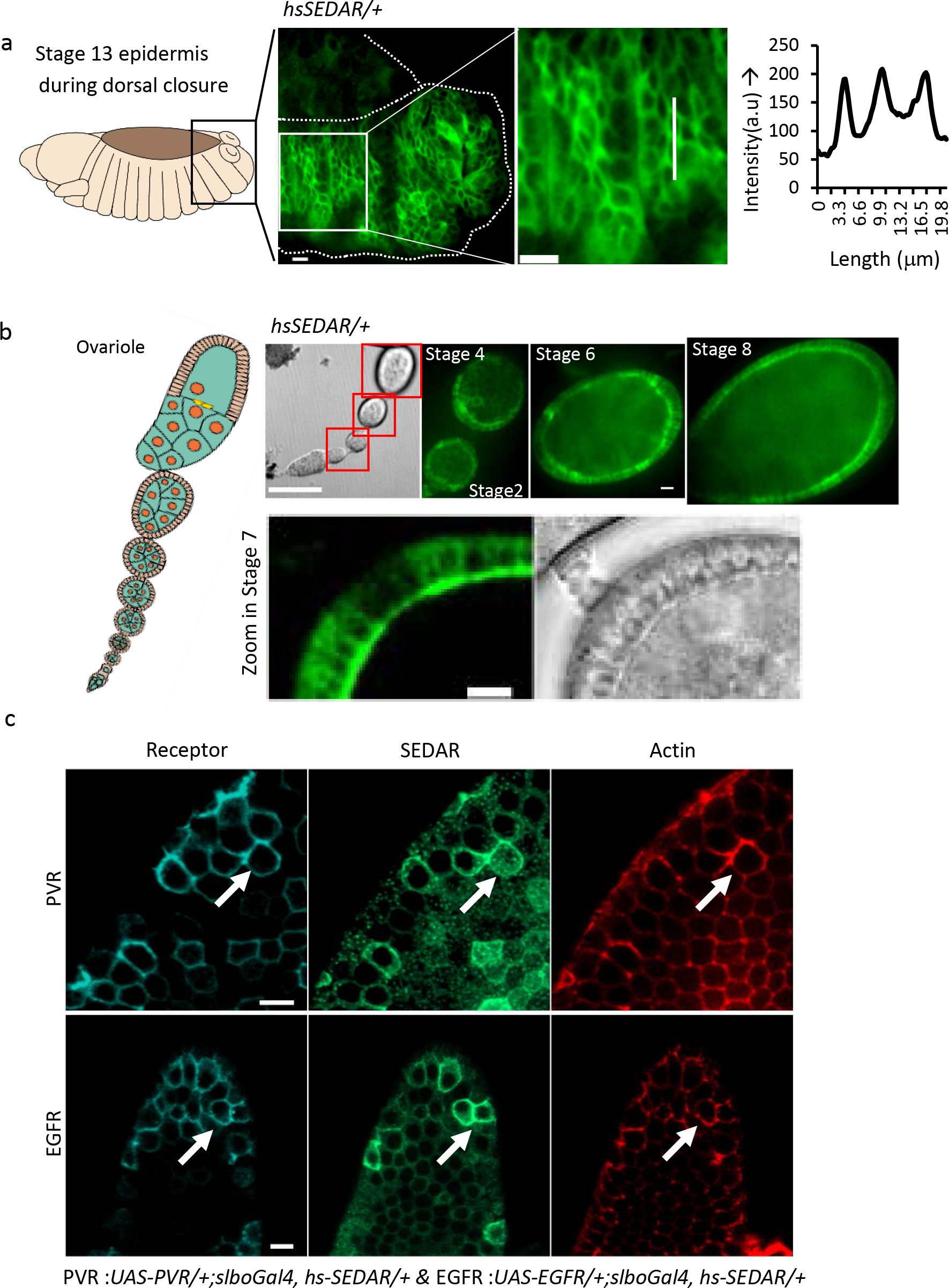
SEDAR detected activated Ras in tissues. (a) schematic of Dorsal closure at stage 13 during embryogenesis, box marked the zoom in region shown in the next panel. Graph of the intensity plot of line drawn in right panel (b) schematic of an ovariole showing the germ line, and ovarioles until stage10. DIC image of egg chamber, scale100μm. Various stages of egg chamber showing SEDAR localized to the apical membrane of follicular epithelium. Scale 10μm. Zoom in of stage 7 egg chamber. (c) Follicular epithelium of stage10 egg chamber expressing slbo-Gal4 and UAS PVR (top panel) or UAS DER (bottom panel) stained with anti PVR or anti DER antibody (cyan), SEDAR (green) and phalloidin (red). Cell with high PVR or DER expression are marked with a white arrow in each panel. Scale 10μm.

During early stages of *Drosophila* oogenesis, Erk is activated in the follicular epithelium. SEDAR localized to the apical plasma membrane of the follicular epithelium in stage 2-7 (Figure 4b). In stage 8, ERK is activated downstream of EGFR in the posterior follicle cells (33). In these cells, SEDAR is localized to the apical plasma membrane of the dorsal follicle cells (Figure 4b).

Overexpression of PVR in the ovariole, under *slboGal4*, leads to high level of PVR expression in few cells of the follicular epithelium, some which in turn activated downstream signals (Figure 4c (15)). In cells overexpressing PVR labeled with total PVR antibody (marked with arrow, Figure 4c (15), SEDAR localized to the plasma membrane whereas no preferential localization was observed in cells not activating PVR within the same tissues (Figure 4c). Similarly, in cells overexpressing EGFR (as observed in EGFR antibody staining, Figure 4c), SEDAR localized to the plasma membrane, whereas diffuse cytoplasmic localization of SEDAR was observed in the neighboring cells not overexpressing EGFR (Figure 4c).

Taken together, this paper demonstrated in 5 different *Drosophila* tissues that SEDAR could detect endogenous activated Ras, particularly downstream of EGFR or PVR at subcellular resolution. Its localization could be monitored in real time in endogenous tissues.

## Discussion

Real-time biosensors that reflect activity of key signaling molecules in the cell have greatly expanded the possibilities for quantitative molecular analysis of dynamic processes such as cell migration but they are limited to analyses in 2D cell culture systems. Observing signaling events in physiological conditions was much needed. In this paper, an approach to study signaling via the small GTPase Ras using highly fluorescent recruitment-based probes in cells is presented. Biosensor, SEDAR, specifically detected Ras activation downstream of growth factor receptors in multiple instances in *Drosophila* epithelial tissue. In cells with irregular membranes such as migrating border cells, SEDAR detected endogenous signaling without any amplification of the signal (overexpression of PVR) indicating that it is more sensitive than antibodies to phosphorylated receptors (15). SEDAR allowed detection of RTK activation *in vivo* in real time.

SEDAR detected signal downstream of the chemotactic RTKs. This detection of endogenous signal provided an evidence of how RTKs could regulate distinct behaviors observed in the clusters. Comparing wild type and guidance-defective clusters, showed that guidance signaling is indeed reflected by SEDAR and detectable by the method presented here. Magnitude of cell intrinsic polarity determined the behavior of the leading cell indicating that the magnitude of subcellular RTK activation was informative to the cell. A higher magnitude of RTK activation at the leading edge could be above the threshold required for Rac activation, which is crucial for cell migration and in turn could promote lamellipodia like large protrusion formation (34–37). Rac is directly downstream of PDGF (38). PVR (PDGF VEGF like receptor) has a high magnitude of cell intrinsic front polarity of SEDAR and Rac activation is high in the leading cell (34, 39). Together this suggests that high magnitude of RTK activation, primarily downstream of PVR can promote formation of large protrusions in the leading edge via Rac. EGFR stimulated small protrusions and it was shown to directly interact with Cdc42 that promotes filopodia like protrusions (40).

Cell extrinsic cluster polarity was informative to guide the forward directed migration of the border cell cluster. It provides a direct evidence for strong local activation and mild global inhibition of RTK signaling (41–44). Single cells that do not have RTK signaling could not occupy the position of the leading cell (45) implying a necessity of high signal in the leading cell. Secondly, cell extrinsic cluster polarity provided an explanation for the polarization of two crucial downstream signals within the cluster i.e. tension in E-cadherin and Rac activation in the front of the border cells cluster (34, 46). Previous evidences fell short of explaining these critical signal gradients essential for migration wherein gradient of Pvf1 in the egg chamber has been difficult to detect and there is no difference in the amount of E-cadherin in the front and back of the cluster. To confirm these results, expression of RTK^DN^led to loss of cell extrinsic cluster polarity and both its features were absent viz, forward migration was severely delayed and it corresponded to similar loss of polarization of tension in E-cadherin (46). Further, cell extrinsic cluster polarity in the direction of migration provided evidence that there is a long-range signal integration of the RTKs over the length scale of the cluster (44). E-cadherin and Rab 11 could mediate long-range communication between border cells along the length of the cluster primarily via cell-cell contacts (46, 47).

In general, the strategy of low-level expression of highly fluorescent recruitment sensors coupled with normalization is useful for many spatially organized signaling events in tissues. Given that the signals probed by the sensors are used repeated in several tissues to bring about a myriad of functions, they could be adapted to any tissue of choice. Most epithelial tissues are very well suited for analyses as they have straight well-defined apical membrane, allowing for easy detection of the signal. With the ratiometric approach, it can be adapted to cells with irregular plasma membrane. Expression under the heat shock promoter can restrict the expression of the sensor temporally thereby allowing for precise analyses at a developmental time of choice. The versatility and universality of the localization based sensor, SEDAR, has a potential to provide many insights into the signaling dynamics of various different RTKs upstream of Ras. Sensors based on designs similar to SEDAR could be adapted to other signaling pathways to visualize them in three-dimensional *in vivo* tissues.

## Acknowledgements

Majority of this work was carried out at Institute of Molecular and Cell Biology, A*STAR, Singapore by the author in the laboratory and guidance of Pernille Rorth. I would like to thank Yusuke Toyoma, Timonthy Saunders, Christopher Amourda, Prabhat Tiwari from Mechanobiology Institute and Cai Yu from Temasek Lifesciences Laboratory for helpful discussions and help with the fly facilities and reagents.

## Materials and Methods

SEDAR is freely available upon request from the author.

### Molecular cloning

For Ras-sensor tests, short or extended Ras-binding domains from Drosophila Raf/polehole coding sequence (supplemental Figure S1) were cloned into pmEGFP10N-1 (kind gift from Jan Ellenberg, EMBL). This was cloned into pCasper-HS and finally into pAttB, transgenic flies made by P-element transformation. For border cell, RAS-sensor, 2 copies of extended RAS binding domains of Drosophila Raf (amino acids 120-307) separated by a 6-glycine linker, were cloned into, and then to pAttB-slbo3.6. pAttB-slbo3.6 was made by inserting a 3.6 kb genomic fragment from upstream of the slbo coding region (including part of the border cell enhancer (Rørth et al 1998) and a long 5’ UTR with several ultra-short ORFs. Primers: Forward-GAAGTGATGCTAGCGGATCCAGCTGCGGCGTTTTATTCTCACGCT and Reverse-CGCAGATCTGTTAACGAATTCTGCAGATTGTTTAGC and poly-adenylation sequences from SV40 into pAttB. Transgenes were insertion into flies with AttP at 86Fb. For 2xslbo-turboRFP, turboRFP [Invitrogen] was sub-cloned downstream of two copies of the slbo enhancer (Rørth 1998) and a minimal promoter for high expression in anterior follicle cells starting at stage 6, transferred to pCasper-AttB and inserted at the VK33 AttP site. AttP flies were obtained from Bloomington.

### Flies and genetics

Fly stocks were obtained from the Bloomington and Vienna Drosophila stock centers. RNAi lines used were GD13502 for *Pvr*, GD43268 for *Egfr* (see Poukkula et al), GD12553 and TRiPGL00336 for *Ras85D*. All UAS-RNAi constructs were expressed under control of actin-flipout-Gal4 (AFG) control, driving robust expression in all somatic cells after heat shock– induced expression (37°C 30 minutes) of FLP recombinase two days prior to dissection and imaging. PVR^DN^, EGFR^DN^, UAS-PVR, UAS-EGFR (see Duckek, Duckek et al) were expressed under slbo-GAL4 control, which drives expression in outer (non-polar) border cells.

For imaging, 2-3 day old females resulting from an outcross, maintained at 25°C, and incubated about 20 hours with wet yeast were used. All in w^1118^mutant background. HS-sensor expression was induced by heat shock at 34°C for 20 minutes and flies were allowed to recover at room temperature for 3 hours prior to imaging.

### Egg chamber dissection and imaging

Ovaries and egg chambers were dissected in dissection medium (Schneider’s medium [Gibco] +5 μg/ml insulin [Sigma]) and stage 9 egg chambers transferred to imaging medium (dissection medium + 2.5% fetal calf serum [Hyclone] + 2 mg/ml trehalose [Fluka] +5 μM methoprene [Sigma] + 1 μg/ml 20-hydroxyecdysone [Sigma] + 50 ng/ml adenosine deaminase [Roche] in poly-lysine-coated imaging chambers [Nunc] at room temperature. Videos were recorded by an inverted confocal microscope [SP5; Leica] with a 63×, 1.2 NA Plan Apochromat water immersion objective, 4x zoom and 2 times line average. The time interval between images was 10 seconds. Egg chambers were aligned by rotating the scan field with the anterior tip of the egg chamber aligned to the left and the image x axis going from this point through the middle of the oocyte (far right). The focal plane with border cells having the furthest front tip was chosen for imaging. After imaging (10 minutes) different focal planes were rechecked to ensure the right focal plane had been used in the whole video. Egg chambers or videos with defects were discarded.

### Image analysis

To generate ratiometric images, GFP and RFP images were first smoothed by Gaussian Blur (sigma 1.0) and then the GFP image was divided by the RFP image (Image J). The front, side and back signals were measured in 1.5μm long and 0.4 μm width lines in ratio-metric images and cytoplasmic signal as a small square box inside the front cell. For each video, 30 to 60 images were measured as long as the leading cell edge or rear cell edge that was being measured was in the optical plane being imaged.

### Immunostaining

After dissection ovaries were fixed in 4% paraformaldehyde for 20 min. Ovaries were washed in buffer [50 mM Tris-HCl (pH 7.4), 150 mM NaCl, 0.05%, 1 mg/mL BSA] for 30 minutes at RT and manually dissociated into single egg chambers by pipetting them using 200ml tip. They were blocked in 5 mg/mL BSA for 30 minutes and incubated overnight at 4 °C with primary antibody 1:200 (16) for total PVR or total EGFR) and subsequently incubated with fluorescently labelled secondary antibody 1:200 (Jackson Immunoresearch Laboratories)+ rhodamine-coupled phalloidin (Molecular Probes).

